# Chronic nicotine exposure is associated with electrophysiological and sympathetic remodeling in the intact rabbit heart

**DOI:** 10.1101/2023.11.23.567754

**Authors:** Amanda Guevara, Charlotte E.R. Smith, Jessica L. Caldwell, Lena Ngo, Lilian R. Mott, I-Ju Lee, inivas Tapa, Zhen Wang, Lianguo Wang, William R. Woodward, G. Andre Ng, Beth A. Habecker, Crystal M. Ripplinger

**Affiliations:** Department of Pharmacology, University of California Davis, Davis, CA, USA; Shantou University Medical College, Shantou, China; Department of Chemical Physiology and Biochemistry, Oregon Health and Science University, Portland, OR, USA; Department of Cardiovascular Sciences, University of Leicester, Leicester, UK; National Institute for Health & Care Research, Leicester Biomedical Research Centre, Leicester, UK; Glenfield Hospital, Leicester, UK; Department of Medicine and Knight Cardiovascular Institute, Oregon Health and Science University, Portland, OR, USA

**Keywords:** Action potential, Arrhythmia, Calcium, Nicotine, Sympathetic activity

## Abstract

Nicotine is the primary addictive component in tobacco products. Through its actions on the heart and autonomic nervous system, nicotine exposure is associated with electrophysiological changes and increased arrhythmia susceptibility. However, the underlying mechanisms are unclear. To address this, we treated rabbits with transdermal nicotine (NIC, 21 mg/day) or control (CT) patches for 28 days prior to performing dual optical mapping of transmembrane potential (RH237) and intracellular Ca^2+^ (Rhod-2 AM) in isolated hearts with intact sympathetic innervation. Sympathetic nerve stimulation (SNS) was performed at the 1^st^ – 3^rd^ thoracic vertebrae, and β-adrenergic responsiveness was additionally evaluated as changes in heart rate (HR) following norepinephrine (NE) perfusion. Baseline *ex vivo* HR and SNS stimulation threshold were increased in NIC vs. CT (*P* = 0.004 and *P* = 0.003 respectively). Action potential duration alternans emerged at longer pacing cycle lengths (PCL) in NIC vs. CT at baseline (*P* = 0.002) and during SNS (*P* = 0.0003), with similar results obtained for Ca^2+^ transient alternans. SNS reduced the PCL at which alternans emerged in CT but not NIC hearts. NIC exposed hearts also tended to have slower and reduced HR responses to NE perfusion. While fibrosis was unaltered, NIC hearts had lower sympathetic nerve density (*P* = 0.03) but no difference in NE content vs. CT. These results suggest both sympathetic hypo-innervation of the myocardium and diminished β-adrenergic responsiveness with NIC. This autonomic remodeling may underlie the increased risk of arrhythmias associated with nicotine exposure, which may be further exacerbated with continued long-term usage.

**NEW & NOTEWORTHY:** Here we show that chronic nicotine exposure was associated with increased heart rate, lower threshold for alternans and reduced sympathetic electrophysiological responses in the intact rabbit heart. We suggest that this was due to the sympathetic hypo-innervation of the myocardium and diminished β- adrenergic responsiveness observed following nicotine treatment. Though these differences did not result in increased arrhythmia propensity in our study, we hypothesize that prolonged nicotine exposure may exacerbate this pro-arrhythmic remodeling.

## INTRODUCTION

Cigarette smoking is the most preventable cause of cardiovascular morbidity and mortality in the United States (1–3). While smoking promotes the progression of coronary artery disease and increases the risk of myocardial infarction (MI), heart failure, ischemia-induced arrhythmias and sudden cardiac death (4–6), short-term secondhand smoke (SHS) exposure alone has been associated with increased incidence of atrial fibrillation and ventricular tachycardia/fibrillation (7). Although the mechanisms for this increased vulnerability to arrhythmias have yet to be fully explored, we have previously shown that the susceptibility to calcium (Ca^2+^) and action potential duration (APD) alternans was increased with SHS exposure (8). At present, however, the contributions of individual cigarette smoke components to the increased emergence of alternans and arrhythmias is unknown. Whereas cigarette usage has declined, sales of electronic (e)-cigarettes increased by 46.6% between 2020 – 2022 (9). Though often considered less harmful than traditional cigarettes, e-cigarettes also contain numerous chemicals including nicotine, tobacco-specific nitrosamines, aldehydes and flavors (10–13). The amount of nicotine present in e-cigarettes is widely variable but, in some cases, has been reported to be higher than an entire pack of conventional cigarettes (14, 15).

Nicotine directly modulates autonomic control of the heart through its binding to nicotinic acetylcholine receptors (nAChR) found within the brain, adrenal medulla, and post ganglionic neurons of the parasympathetic and sympathetic cardiac nerves (16). In studies using labeled norepinephrine (NE), rats given nicotine for 10 days had a higher NE but lower acetylcholine outflow in the hippocampus (16), indicating that the overall net effect from the central nervous system is sympathetically driven and that chronic nicotine exposure could lead to sympathetic hyperactivity. Similarly, acute exposure to e-cigarettes containing nicotine has been linked to reduced heart rate variability (HRV) (17), which is typically associated with increased sympathetic activity and risk of sudden cardiac death (18, 19). Thus, it is possible that long-term nicotine use results in autonomic dysregulation that may play a role in promoting ventricular arrhythmias.

In addition to its actions on the autonomic nervous system, nicotine also directly affects the myocardium, likely through its binding to non-neuronal nAChRs found on ventricular cardiac myocytes (20). Acute nicotine administration has been shown to directly block the membrane stabilizing current I_k1_ in isolated myocytes (21, 22) which we have previously demonstrated leads to the generation of ectopic beats in the intact heart (23). Exposure to nicotine is also associated with increased myocardial fibrosis (24–26), which may play a role in creating a heterogeneous substrate that can foster ectopic activity and/or reentrant arrhythmias (27). In line with these findings, *in-vivo* administration of nicotine was reported to result in atrial and ventricular arrhythmias in a canine model of MI (28, 29), with meta-analysis studies also identifying increased risk of rhythm disorders in patients on nicotine replacement therapies (30, 31). As such, at the level of the heart, nicotine exposure is associated with myocardial remodeling alongside alterations in autonomic regulation that appear to be synergistically proarrhythmic. However, despite the importance of these findings, the electrophysiological mechanisms underlying increased arrhythmia susceptibility due to nicotine exposure are unclear. Therefore, the goal of this study was to investigate the effect of chronic nicotine exposure on cardiac electrophysiology, Ca^2+^ handling, structure, and sympathetic responsiveness in the intact heart to elucidate potential mechanisms of increased arrhythmia risk.

## MATERIALS AND METHODS

### Ethical Approval

All procedures were approved by the Animal Care and Use Committee of the University of California, Davis and were conducted according to the Guide for the Care and Use of Laboratory Animals published by the National Institutes of Health (NIH publication no. 85-23, revised 2011).

### Nicotine Exposure

Male (N = 10) and female (N =15) New Zealand White rabbits (4 – 8 months old, 3.0 – 3.5 kg) were obtained from Charles River and singly housed with *ad libitum* access to food and water. Rabbits were randomly assigned to control (CT) or nicotine (NIC) groups with exposure carried out in 4 similarly sized paired cohorts. Exposure to nicotine occurred via transdermal patches (Nicoderm CQ, 21 mg/day) placed on the ears, with surgical tape used for control rabbits. Transdermal patches and surgical tape were changed daily Monday – Friday with a total exposure duration of 28 days.

### Measurement of Serum Cotinine Levels

To confirm delivery of nicotine, levels of the circulating nicotine metabolite cotinine were assessed using blood drawn from the marginal ear vein of NIC and CT rabbits on day 0, 14 and 28 of exposure. Blood samples were allowed to clot overnight at 2 – 8 °C prior to centrifugation at 1000 *g* (3000 rpm) for 15 minutes. Serum was then removed and stored at -80 °C. To assess serum cotinine levels, samples were sent overnight on dry ice to the Tobacco and Carcinogen Biomarkers Core at the University of California San Francisco for analysis by liquid chromatography-tandem mass spectrometry (LCMSM) as previously described (32).

### Innervated Whole-Heart Langendorff Perfusion

At 28 days of exposure, rabbits were euthanized via intravenous injection of pentobarbital sodium (100 mg/kg) following administration of 1000 IU heparin (Fresenius Kabi USA, IL). Hearts were surgically extracted with the thoracic spinal column (T1 – T12) attached to ensure intact sympathetic innervation as in previous studies (33, 34). Following extraction, the isolated preparation was flushed with ice cold cardioplegia (composition in mmol/L: NaCl 110, CaCl_2_ 1.2, KCl 16, MgCl_2_ 16, and NaHCO_3_ 10) via the descending aorta. The preparation was then retrograde perfused and submerged in Tyrode’s solution at 37 ± 0.5 °C (composition in mmol/L: NaCl 128.2, CaCl_2_ 1.3, KCl 4.7, MgCl_2_ 1.05, NaH_2_PO4 1.19, NaHCO_3_ 20 and glucose 11.1). The excitation-contraction uncoupler (blebbistatin, 10 – 20 µM, R&D Systems INC, Minneapolis, MN, USA) (35) and a skeletal muscle paralytic (vecuronium bromide, 6 µM, Cayman Chemical Company INC, Ann Arbor, MI, USA) were added to the circulating perfusate to eliminate motion during optical recordings. Two needle electrodes were positioned in the thoracic cavity and a ground was placed in the bottom of the perfusion dish to obtain a continuous ECG recording. Perfusion pressure was maintained at 60 – 80 mmHg by adjusting perfusion flow rate (80 – 100 mL/min).

### Optical Mapping

Dual optical mapping was performed as previously described (33, 36–38). Briefly, preparations were loaded with Ca^2+^ (Rhod-2 AM, 50 µL of 1 mg/mL in DMSO + 10% pluronic acid, Biotium, Hayward, CA) and voltage (V_m_)-sensitive dyes (RH237, 50 µL of 1 mg/mL in DMSO, Biotium, Hayward, CA) through the coronary perfusion. A catheter electrode was inserted up the spinal canal to T1 -T3 in order to stimulate the pre-ganglionic sympathetic nerves. A pacing electrode was placed on the center of the left ventricle for epicardial ventricular pacing. The anterior epicardial surface was illuminated with LED light sources at 530 nm bandpass filtered from 511 to 551 nm (LEX-2, SciMedia, Costa Mesa, CA, USA) and focused directly on the surface of the heart. Emitted florescence was captured utilizing a THT-macroscope (SciMedia) and split by a dichroic mirror set at 630 nm (Omega, Brattleboro, VT, USA). RH237 signals were longpass filtered at 700 nm, while Rhod-2 AM signals were bandpass filtered from 574 to 606 nm. Signals were recorded with two CMOS cameras (MiCam Ultima-L, SciMedia) at a sampling rate of 1 kHz and a 100 x 100 pixel resolution, with a field of view of 3.1 x 3.1 cm and a spatial resolution of 0.31 mm/pixel.

### Experimental Protocol

*Ex vivo* heart rate (HR) was monitored prior to the addition of blebbistatin and throughout experiments via continuous ECG recording in LabChart (ADInstruments INC, Colorado Springs, CO, USA). After the addition of blebbistatin, sympathetic nerve stimulation (SNS) thresholds were determined as previously described (36). Specifically, sympathetic nerve fibers were initially stimulated at 0.5 Hz, 7 V for 5 seconds. Stimulation frequency was then increased in 0.5 Hz increments at a constant voltage of 7 V until a > 15% increase in HR was observed. Sympathetic nerve stimulation thresholds were then noted and hearts subsequently stimulated at 5 Hz higher than their threshold value. Baseline Ca^2+^, V_m_, and HR recordings were made with the absence or presence of 13 seconds of SNS. Continuous ventricular pacing was then performed via the epicardial left ventricle (LV) at decreasing pacing cycle lengths (PCL) from 300 ms until hearts lost capture. Finally, hearts were challenged with NE (500 nM) in the perfusate to evaluate β- adrenergic responsiveness.

### Optical Mapping Data Analysis

Optical data was analyzed using both Optiq (Cairn Research, UK) and ElectroMap (39) software as described previously (33, 40, 41). Briefly, APD (APD_80_) or Ca^2+^ transient duration (CaTD_80_) was calculated as 80% repolarization minus activation time (with activation time calculated at 50% of peak amplitude). Mean APD_80_ and CaTD_80_ for each heart were calculated from the entire mapping field of view. CaTD and APD alternans thresholds were determined using spectral methods as previously described (42, 43) with the threshold defined as the slowest PCL at which significant alternans emerged (minimum spectral magnitude ≥ 2).

Due to SNS electrical artifacts on the ECG, data pertaining to changes in HR during SNS were analyzed from optical recordings using ElectroMap. All other HR measurements were obtained from continuous ECG recordings and analyzed using LabChart (ADInstruments INC, Colorado Springs, CO, USA). *Ex vivo* baseline HR was measured 1 minute prior to the addition of blebbistatin and averaged over 1 minute. For HR changes in response to NE, HR was measured 1 minute prior to the addition of NE (averaged over 10s) and 6 minutes after NE perfusion. To quantify incidence and severity of arrhythmia, ECG recordings were scored according to previous studies (36, 44, 45). Briefly, 0 = no ectopic beats, 1 = premature ventricular contraction (PVC), 2 = bigeminy or salvos (2 – 4 consecutive beats), 3 = ventricular tachycardia (> 5 consecutive, monomorphic beats), or 4 = ventricular fibrillation (> 5 consecutive, polymorphic beats). Arrhythmia scores were determined as the most severe arrhythmia observed during the entire experimental protocol.

### Tissue harvesting

After completion of optical mapping experiments, a 2 mm thick short-axis section was cut from the middle of the LV and sliced into wedges for multiple biochemical experiments. For immunohistochemical and histological staining, LV tissue wedges were fixed in 4% paraformaldehyde for 1.5 hours, washed with phosphate buffered saline (PBS), and placed in 30% sucrose in PBS overnight. Tissue was then embedded in Optimal Cutting Temperature (OCT) medium, flash frozen, and stored at -80 °C. For high performance liquid chromatography (HPLC), LV and right atrial (RA) tissue were flash frozen and stored at -80 °C.

### Assessment of Fibrosis

To assess fibrosis, tissue sectioning and Masson’s Trichrome staining was performed by Acepix Biosciences Inc (Hayward, CA, USA). Tissue was sectioned into 10 µm thick slices at a step depth of 50 – 100 µm, thaw-mounted on positively charged slides, stained and stored at -80 °C. Imaging was performed on an upright Nikon Eclipse Ni microscope at 20 x magnification with white light excitation. Fibrosis area was color-thresholded (blue) and divided by total tissue area. All images were analyzed by two blinded users and the generated values averaged.

### Assessment of Sympathetic Nerve Density

To assess sympathetic nerve density, tyrosine hydroxylase (TH) labeling was performed as in previous studies (36, 41, 46). Briefly, slides (N = 6 slices/group) were rehydrated with PBS and incubated in sodium borohydride (10 mg/mL) in order to reduce background auto-florescence. Slides were then blocked with 2% bovine serum albumin (BSA, Sigma) and 0.3% Triton X-100 (Sigma) in PBS (BSA/PBS-T) for 1 hour prior to washing with PBS and incubation with primary anti-TH rabbit polyclonal antibody (EMD Millipore) at 1:300 BSA/PBS-T overnight at room temperature. Slides were subsequently rinsed in PBS and incubated in Alexa Fluor 488 conjugated goat anti-rabbit secondary antibody (1:500; Invitrogen) for 1.5 hours at room temperature. After incubation, slides were rinsed with PBS, briefly dipped in MilliQ water prior to incubation in a 10 mM copper sulfate/50 mM ammonium acetate solution to reduce autofluorescence (3 x 10 minutes). Finally, slides were briefly re-dipped in MilliQ water and a 1:1 glycerol/PBS solution added to mount a coverslip. Imaging was performed on an upright Nikon Eclipse Ni microscope at 10 x magnification with a FITC filter (Ex/Em: 495/519 nm). Four images per tissue slice were taken and thresholded in ImageJ to determine TH+ sympathetic nerve fiber area and total tissue area. Percent fiber density was defined as the ratio of TH+ fiber area to total tissue area. As for fibrosis, all images were analyzed by two blinded users and values averaged between users.

### Tissue Norepinephrine Content

NE content was measured via HPLC as described previously (36, 47). Frozen tissue was homogenized at 25 °C in perchloric acid (PCA, 0.1 M) containing the internal standard dihydroxybenzylamine (DHBA, 0.25 µM) to correct for NE sample recovery. Catecholamines were purified from a 100 µL aliquot of homogenate using alumina extraction. NE was reabsorbed from alumina by using PCA (150 µL, 0.1 M) and then measured by C18 reversed-phase HPLC with electrochemical detection (ESA, Coulchem III) and a detection limit of ∼ 0.05 pmol and > 60% recovery from alumina extraction.

### Statistics

Data are expressed as mean ± standard deviation (SD) for N animals. Due to the technical challenges of sequential *in vivo* blood collection and some *ex vivo* experimental procedures, sample sizes and sex distribution are unequal for some parameters. Likewise, in some cases the *ex vivo* innervated heart preparations did not capture at a particular pacing frequency, which impacted sample size for some rate-matched parameters. N are specified in each figure legend and data from both sexes pooled, with O and X symbols indicating female and male sex, respectively (48, 49). Statistical analysis was performed using GraphPad Prism 9. Normality was tested using the Shapiro-Wilk test and significance was assessed using t-test, two-way repeated measures ANOVA, two-way ANOVA with mixed effects analysis or Mann-Whitney test as appropriate and specified in the figure legend. Statistical significance was attained when *P* < 0.05.

## RESULTS

### Baseline Measurements Following Nicotine Administration

To confirm nicotine delivery, cotinine was measured in serum blood samples. As expected, CT rabbits had no detectable cotinine at all time points (**Fig. 1A**). In NIC rabbits, there was no detectable cotinine at day 0, but 455.3 ± 374.0 ng/mL and 694.6 ± 376.4 ng/mL of cotinine at 2 and 4 weeks of exposure, respectively. To determine the impact of nicotine on *ex vivo* HR; baseline HR measurements were obtained from the innervated perfused heart preparations prior to the addition of blebbistatin and dyes. HR was increased by 23.8 ± 32.7 % in NIC hearts compared to CT (**Fig. 1B**, 206.3 ± 24.5 BPM vs. 166.6 ± 38.8 BPM, *P* = 0.004). SNS stimulation thresholds were also increased in NIC hearts (**Fig. 1C**, 7.3 ± 5.3 Hz vs. 2.1 ± 1.0 Hz, *P* = 0.003).

**Figure 1:**
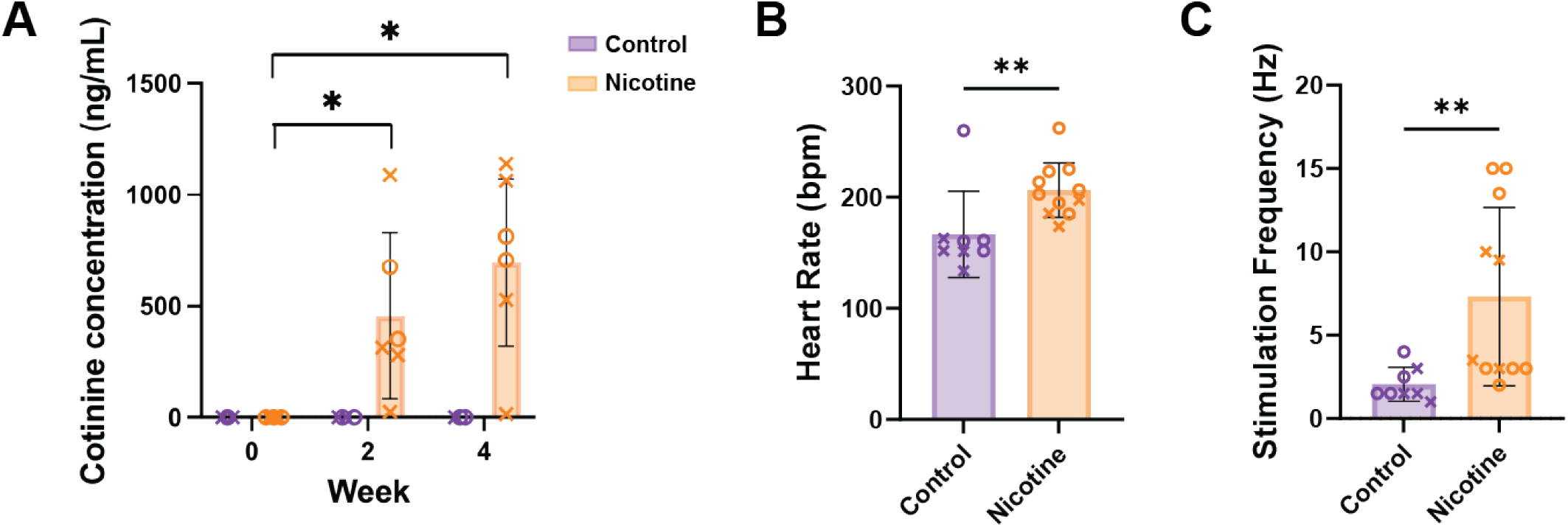
Cotinine concentration, heart rate, and sympathetic nerve stimulation threshold in control and nicotine-exposed hearts. (A) Mean cotinine concentration at 0, 2, and 4 weeks of nicotine exposure. N=6-10/group. (B) *Ex vivo* baseline heart rate. Measurements taken prior to the addition of dyes and blebbistatin. (C) Stimulation frequency threshold for sympathetic nerve stimulation. Spinal cord was stimulated at T1 - T2 with constant voltage and frequency was increased 0.5 Hz increments until a >15% increase in heart rate was observed. N=8-11/group. Data are mean ± SD. * *P* < 0.05, ** *P* < 0.01, by two-way ANOVA with mixed-effects analysis (A) or by Mann-Whitney test (B, C). X denotes males, O denotes females.

### Electrophysiological Responses to SNS

To determine the effect of SNS on electrophysiological parameters, HR, APD_80_, and CaTD_80_, were measured immediately prior to and at 13 seconds of SNS. SNS resulted in comparable increases in HR in both CT and NIC hearts (**Fig 2B**, 41.1 ± 45.52% vs. 29.2 ± 32.66%, *P* = 0.51). Similarly, SNS shortened APD_80_ to a similar degree in both CT and NIC hearts (**Fig. 2C-D**). Interestingly, SNS reduced CaTD_80_ compared to baseline values in CT but not NIC hearts (**Fig. 2D**, *P* = 0.018), however the % change in CaTD_80_ with SNS was not different between groups (**Fig. 2E**, *P* = 0.411).

**Figure 2:**
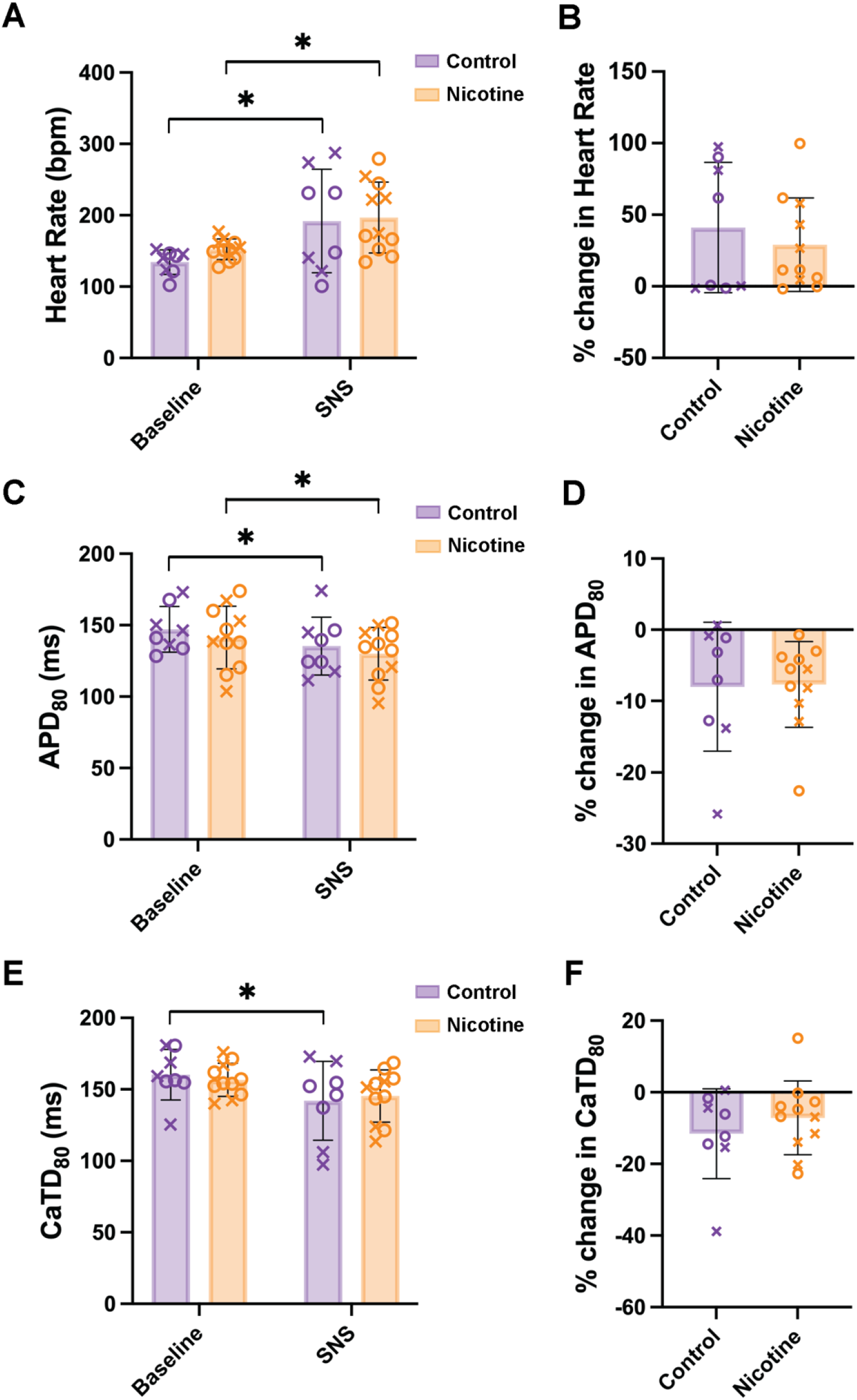
Effect of sympathetic nerve stimulation (SNS) on control and nicotine-exposed hearts. Heart rate (A-B), action potential duration (APD_80_, C-D), and calcium transient duration (CaTD_80_, E-F) changes with SNS. N=8-11/group. Data are mean ± SD. * *P* < 0.05, by two-way ANOVA with repeated measures (A, C, E), Mann-Whitney test (B) or two-tailed, unpaired t test (D, F). X denotes males, O denotes females.

### APD and CaTD Alternans

To evaluate alternans susceptibility, hearts were paced at progressively shorter PCLs. As sympathetic stimulation may decrease alternans PCL threshold due to accelerated Ca^2+^ handling (33, 42, 50), this was firstly performed without and subsequently with SNS. In both the absence and presence of SNS, APD alternans emerged at longer PCLs in NIC compared to CT hearts (**Fig. 3A**, 238.8 ± 15.5 ms vs. 197.5 ± 22.5 ms, *P* = 0.0022 and 230.0 ± 22.7 ms vs. 180.0 ± 28.3 ms, *P* = 0.0003 respectively). While SNS was associated with alternans emerging at shorter PCLs compared to baseline in CT (**Fig. 3A**, *P* = 0.0194), no difference in APD alternans threshold was observed with SNS in NIC hearts (**Fig. 3A**, *P* = 0.2896). Similar results were obtained for CaTD alternans (**Fig. 3B**), whereby alternans emerged at shorter PCLs in CT animals with SNS vs. baseline (*P* = 0.0013), and the threshold PCL for alternans in the presence of SNS was lower in NIC vs. CT (*P* = 0.003).

**Figure 3:**
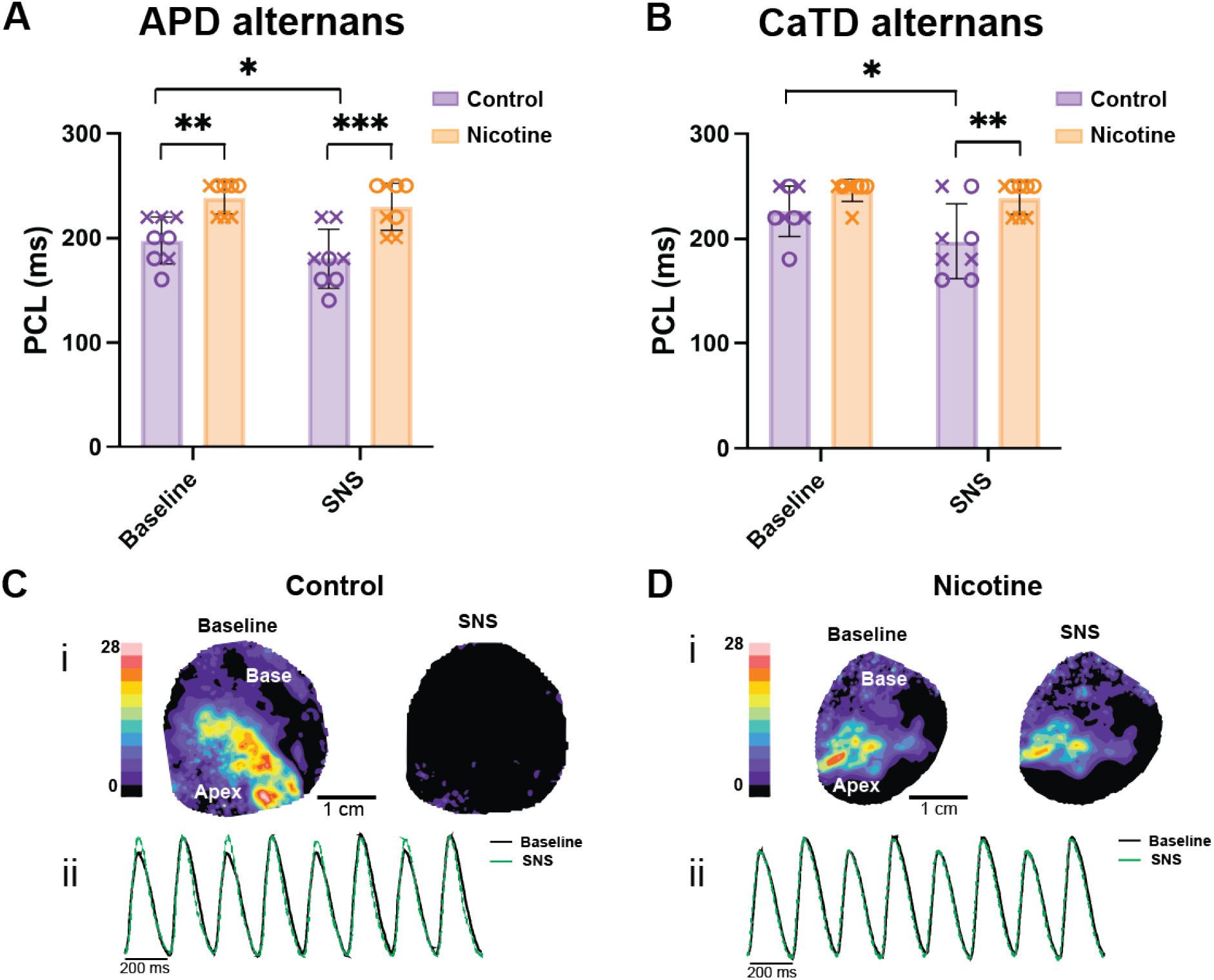
Alternans threshold and magnitude in control and nicotine-exposed hearts. (A) Pacing cycle length (PCL) at which APD alternans emerged at baseline and with sympathetic nerve stimulation (SNS). (B) PCL at which Ca^2+^ alternans emerged. (C-D), representative contour maps and corresponding CaTD traces demonstrating Ca^2+^ alternans magnitude at baseline and with SNS (C, control; D, nicotine). N = 8/group. Data are mean ± SD. * *P* < 0.05, ** *P* < 0.01, *** *P* < 0.001, by two-way ANOVA with repeated measures. X denotes males, O denotes females.

### β-Adrenergic Responsiveness

Electrophysiological differences in response to SNS may be due to altered nerve function (e.g., NE release), altered β-adrenergic responsiveness of cardiomyocytes, or a combination of both. To specifically assess β-adrenergic responses of the heart, 500 nM NE was added the perfusate. HR prior to the addition of NE (time 0, t0) up to 6 minutes after the addition of NE is shown in **Fig 4A**. To account for differences in baseline HR, the % change in HR over 6 minutes of NE perfusion was also calculated (**Fig. 4B**). CT hearts tended to have a larger relative change in HR from initial values compared to NIC (*P* = 0.0591). Both groups had faster HRs at 2 and 3 min after NE compared to t0 (**Fig. 4C**), but only CT hearts showed significant elevation at 1 min (*P* = 0.0005), suggesting a potentially slower HR response in NIC hearts. Maximal HRs after the addition of NE were not different between groups (CT: 216.4 ± 51.4 BPM vs. NIC: 205.2 ± 45.5 BPM, *P* = 0.86).

**Figure 4:**
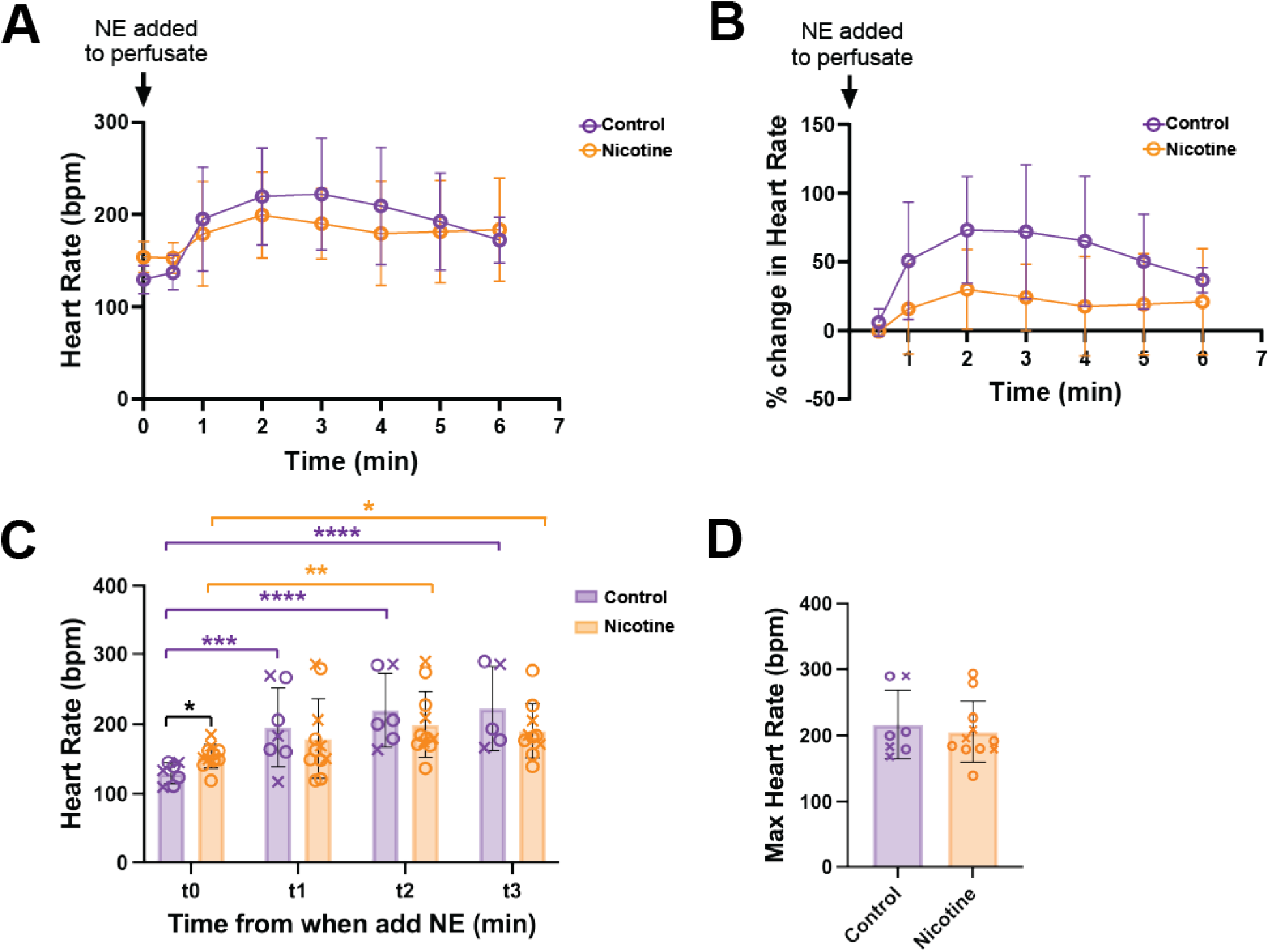
Heart rate changes with norepinephrine (NE) in control and nicotine-exposed hearts. (A-B) Heart rate changes over time after the addition of 500nM NE to the perfusate. N=7-11/group. (C) Heart rate measurements after the addition of NE at 1 minute (t1, N=7-11/group), 2 minutes (t2, N=6-11/group), and 3 minutes (t3, N=5-10/group). (D) Maximum heart rate after NE added, N=7-11/group. Data are mean ± SD. * *P* < 0.05, ** *P* < 0.01, *** *P* < 0.001, **** *P* < 0.0001, by two-way ANOVA with mixed-effects analysis (A, B, C) or Mann-Whitney test (D). X denotes males, O denotes females.

### Arrhythmia Susceptibility

Arrhythmia susceptibility was measured from pseudo ECG recordings obtained from needle electrodes inserted into the posterior thoracic cavity (hence atypical ECG morphology). PVCs and bigeminy/salvos were observed in both CT and NIC hearts with no difference in arrhythmia score determined between groups (**Fig. 5**, CT: 0.88 ± 0.83 vs. NIC: 1.5 ± 0.71, *P* = 0.1354).

**Figure 5:**
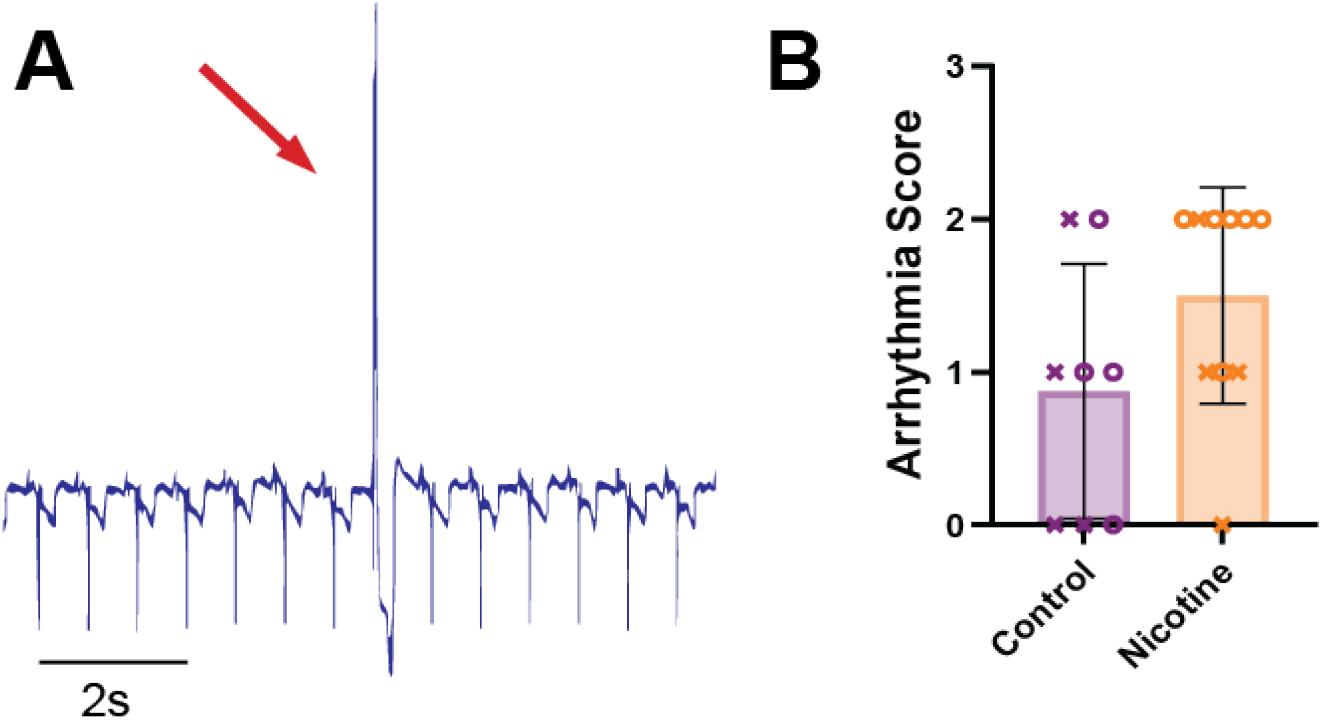
Arrhythmia susceptibility in control and nicotine-exposed hearts. Pseudo-ECG recording demonstrating arrhythmias during sinus rhythm. Recordings were measured from the posterior thoracic cavity in the bath. (A) example of PVC. (B) Average arrhythmia scores per group. N=8- 10/group, Data are mean ± SD. Significance tested by Mann-Whitney test. X denotes males, O denotes females.

### Fibrosis, Sympathetic Nerve Density, and Norepinephrine Content

Chronic nicotine administration has been previously associated with increased myocardial fibrosis (24–26). Therefore, fibrotic density in CT and NIC hearts was evaluated from Masson’s trichrome images. However, no difference in fibrotic density was found (**Fig. 6A-B**, 21.78 ± 3.68% vs. 19.30 ± 3.68%, *P* = 0.53). To assess whether changes in sympathetic nerve density were responsible for altered SNS responses, the % TH+ fiber density was quantified from LV tissue. CT hearts had a higher % area of TH+ compared to NIC hearts (**Fig. 6C-D**, 10.18 ± 2.45 % vs. 7.55 ± 0.91%, *P* = 0.0339). NE content was also measured in the LV and RA in CT and NIC hearts (**Fig. 6E**). In line with greater sympathetic nerve density in the atrium vs. ventricle (51), we observed increased NE content in the RA vs. LV (*P* = 0.0037, two-way ANOVA main effect). Pairwise, we found increased NE content in the RA vs LV of NIC hearts (*P* = 0.0194), and a tendency for this in CT (*P* = 0.1264). Interestingly, despite the lower ventricular nerve density, we saw no difference in NE content in the NIC LV compared to CT (29.2 ± 9.5 pmol/mg vs. 25.6 ± 8.4 pmol/mg, *P* = 0.7431), though a tendency for increased NE was observed in the RA (44.6 ± 12.1 pmol/mg vs. 34.8 ± 9.3 pmol/mg, *P* = 0.1970).

**Figure 6:**
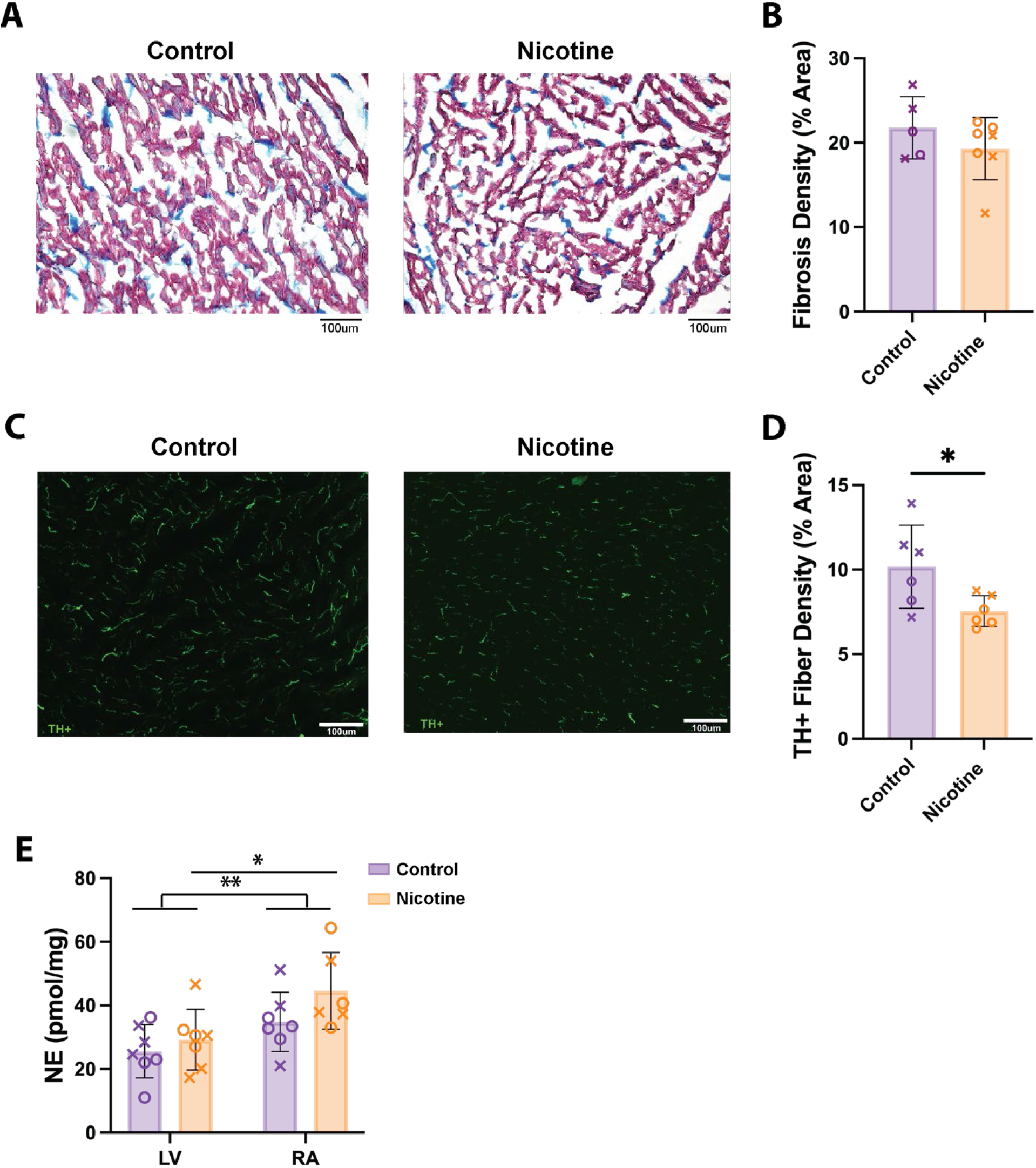
Fibrosis, sympathetic nerve density, and norepinephrine (NE) content in control and nicotine-exposed hearts. (A) Representative images of Masson’s Trichrome staining; fibrosis indicated in blue and myocardial tissue in red. (B) Average fibrosis density in left ventricle. N =5–7/group. (C) Representative immunofluorescence images of tyrosine hydroxylase (TH) labeling. (D) Sympathetic nerve fiber density in left ventricle. N=6/group. (E) NE content of left ventricle (LV) and right atria (RA). N=7/group. Data are mean ± SD. * *P* < 0.05, ** *P* < 0.01, by Mann-Whitney test (B), two-tailed, unpaired t test (D) or two-way ANOVA with repeated measures (E). Scale bar = 100 µm. X denotes males, O denotes females.

## DISCUSSION

The objective of this study was to investigate the effects of chronic nicotine exposure on cardiac sympathetic responses, electrophysiological and structural remodeling, and potential arrhythmogenic effects. To our knowledge, this study is the first to employ the innervated rabbit heart model to evaluate sympathetic responses in hearts chronically exposed to nicotine. Here, we show that NIC hearts had: (1) higher *ex vivo* baseline HRs; (2) elevated SNS thresholds and reduced sympathetic nerve density; (3) increased susceptibility to Ca^2+^ and APD alternans that does not improve with SNS; (4) altered ß- adrenergic responsiveness; and (5) no change in arrhythmia propensity. Taken together, these results suggest that chronic nicotine exposure leads to sympathetic remodeling and altered cardiac electrophysiological responses, but that these changes are not necessarily arrhythmogenic following 28 days of exposure.

### Chronic Nicotine Exposure Increases Ex Vivo Baseline Heart Rate

Perfusion of a low dose nicotine has previously been shown to result in an acute increase in HR in isolated rabbit hearts (52). Additionally, it has been shown that chronic nicotine administration led to a reduction in HRV that suggests a shift to sympathetic dominance and decreased vagal tone (17, 53). Here, we show that the hearts of rabbits exposed to nicotine for 28 days had elevated baseline *ex vivo* HR in the innervated heart preparation (**Fig. 1B**). Since the sympathetic innervation to the heart remained intact in this preparation, this result may suggest an elevation in basal sympathetic nerve activity under resting conditions (without electrical stimulation) following nicotine exposure. Alternatively, the increase in baseline HR may be due to functional and structural alterations within the SA node itself, which may represent an important area for future study.

### Electrophysiological Responses to SNS After Chronic Nicotine Exposure

While we found that NIC hearts had significantly increased electrical SNS thresholds (measured by the stimulation frequency required to produce a 15% increase in HR, **Fig. 1C**), similar changes in APD, CaTD, and HR were observed with SNS at supra-threshold frequency stimulation (**Fig. 2**). The elevated SNS thresholds could be due to many factors, including changes in sympathetic nerve density or function, altered β-adrenergic responsiveness, or structural remodeling of the myocardium. Chronic nicotine administration is associated with increased atrial fibrosis (24–26); however, we found no difference in ventricular fibrosis between groups in this study (**Fig. 6A**), indicating potential chamber differences in the fibrotic remodeling induced by nicotine. We did observe a reduction in sympathetic nerve density in NIC hearts (**Fig. 6C**) that could explain the blunted responsiveness to SNS. However, despite the decrease in nerve density, we observed no difference in NE content in the RA and LV of NIC hearts compared to CT (**Fig. 6E**). Typically, NE content is associated with local sympathetic nerve density whereby more nerves = more NE. However, sustained activation of nicotinic receptors in postganglionic sympathetic neurons stimulates the activity of TH, increasing NE synthesis and allowing the neuron to maintain elevated transmitter release during extended periods of activation (54, 55). Related data were obtained in human and isolated rabbit heart studies showing that acute nicotine administration leads to an increase in NE release (56–58). Finally, studies in rats with chronic administration of nicotine showed an increase in cardiac NE turnover, but no change in cardiac NE content (59). Thus, our data connecting decreased nerve density with normal NE content is consistent with studies showing that nicotine stimulates NE synthesis in sympathetic neurons and regulates all aspects of noradrenergic transmission independent of nerve density.

### Chronic Nicotine Exposure Increases the Susceptibility for APD and CaT Alternans

We found that NIC hearts had APD and CaT alternans that emerged at slower PCLs compared to CTL (**Fig. 3A & 3B**), indicating increased alternans susceptibility. Our result is corroborated by a study in post-MI dogs that showed an increase in depolarizing and repolarizing alternans after nicotine infusion (28) and our previous work showing an increase in the magnitude of APD and CaT alternans after SHS exposure (8). While others have shown increased monophasic APD_50_ alternans with SNS (60), we expected that the PCL at which alternans emerges would decrease with SNS given β-adrenergic stimulation accelerates Ca^2+^ cycling (42, 50). Indeed, we found that SNS decreased the alternans threshold PCL in CTL hearts but did not improve alternans thresholds in NIC hearts (**Fig. 3A & 3B**). This result could be explained by decreased sympathetic responsiveness (potentially due to diminished sympathetic innervation) that may be particularly evident at faster PCLs. It is known that APD and Ca^2+^ alternans are strongly associated with increased risk for ventricular arrhythmias (42, 61, 62); however, we did not observe an increase in arrhythmia susceptibility in NIC hearts in this study. It is possible that increased arrhythmogenic activity may be observed with longer duration nicotine exposure.

### β-Adrenergic Responsiveness to NE after Chronic Nicotine Exposure

Because SNS may result in variable NE release throughout the heart (due to variations in nerve density and function), NE was directly added to the perfusate to assess β-adrenergic responsiveness of the myocardium. NIC hearts tended to have smaller and slower changes in HR compared to CT (**Fig. 4**). This may be a result of elevated HR in NIC hearts at time 0 (therefore any HR increases may be more modest) but could also suggest a somewhat blunted β-adrenergic response. This blunted β-adrenergic response could be due to loss of coupling between nerves and β-adrenergic receptors (63) or β-adrenergic receptor down regulation, as prior studies previously showed decreased β-adrenergic receptor density and reduced catecholamine response in habitual smokers vs. non-smokers (64). This is also consistent with the sustained increase in cardiac NE release following nicotine exposure that has been observed in human and animal studies (56–58). As our study focused on changes in functional parameters due to chronic nicotine exposure, future studies would need to be performed in order to further elucidate these findings.

### Limitations

With 28 days nicotine exposure we saw a change in some, but not all, electrophysiological parameters. It is possible that with longer exposure duration, there may be more profound changes in electrophysiology, Ca^2+^ handling, and arrhythmia susceptibility. This could form part of a follow-up study which could consider whether nicotine-induced remodeling is reversible and may be important considering the use of nicotine patches as a smoking withdrawal strategy. We specifically focused on ventricular action potentials, Ca^2+^ transients, and nerve density in this study and therefore did not assess corresponding atrial parameters which may be more severely impacted by nicotine exposure due to greater innervation of the atria. While we included both sexes in our study, we were insufficiently powered to assess sex differences in the context of chronic nicotine exposure, and this remains an important area for future study as sex differences in repolarization following e-cigarette use have recently been reported (65).

### Conclusions

Using the innervated rabbit heart model, we determined the effects of chronic nicotine exposure on cardiac electrophysiology, Ca^2+^ handling, and sympathetic remodeling and responses. We found that chronic nicotine exposure results in reduced response to SNS, as indicated by elevated electrical stimulation thresholds. Following nicotine exposure, SNS also failed to slow the emergence of cardiac alternans, which may be pro-arrhythmic. These results are likely due to a combination of sympathetic hypo-innervation of the myocardium and diminished β-adrenergic responsiveness following nicotine exposure. Taken together, our data indicate that chronic nicotine exposure for just 28 days results in potentially detrimental sympathetic and electrophysiological remodeling that we suggest may be further exacerbated with continued longer-term usage.

## DATA AVAILABILITY

Full datasets are available from the corresponding author upon reasonable request.

## GRANTS

This study was funded by the University of California Tobacco Related Disease Research Program (T29IP0365C), the National Institutes of Health (R01 HL111600, R01 HL093056, and T32 GM144303), and the National Natural Science Foundation of China (NSFC, 82200346). G.A.N. has been supported by a British Heart Foundation Programme Grant (RG/17/3/32,774) and the Medical Research Council Biomedical Catalyst Developmental Pathway Funding Scheme (MR/S037306/1).

## AUTHOR CONTRIBUTIONS

A.G., G.A.N., B.A.H., and C.M.R. conceived and designed research; A.G., C.E.R.S., J.L.C., L.N., L.M., I-J.L., S.T., Z.W., L.W., and W.R.W. performed experiments; A.G., C.E.R.S., L.N., W.R.W., B.A.H., and C.M.R. analyzed data and interpreted results of experiments; A.G., C.E.R.S., and C.M.R. prepared figures and drafted the manuscript; all authors edited and revised the manuscript and approved the final version of manuscript.

